# Reduced complexity of brain and behavior due to MeCP2 disruption and excessive inhibition

**DOI:** 10.1101/2020.12.29.424632

**Authors:** Jingwen Li, Patrick A. Kells, Shree Hari Gautam, Woodrow L. Shew

**Affiliations:** University of Arkansas, UA Integrative Systems Neuroscience Group, Department of Physics, Fayetteville, 72701, USA

## Abstract

Rett syndrome (RTT) is a devastating neurodevelopmental disorder, caused by disruptions to the MECP2 gene, and resulting in severe cognitive and motor impairment. Previous work strongly suggests that healthy MECP2 function is required to have a normal balance between excitatory and inhibitory neurons. However, the details of how neural circuit dynamics and motor function are disrupted remain unclear. How might imbalanced E/I cause problems for motor function in RTT? We addressed this question in the motor cortex of awake, freely behaving rats, comparing normal rats with a transgenic rat model of RTT. We recorded single-unit spiking activity while simultaneously recording body movement of the rats. We found that RTT rats tend to have excessive synchrony among neurons in the motor cortex and less complex body movements. Importantly, greater synchrony was correlated with greater stereotypy of relationships between neurons and body movements. To further test how our observations were related to an E/I imbalance, we pharmacologically altered inhibitory synaptic interactions. We were able to recapitulate many of the phenomena we found in MECP2-deficient rats by enhancing inhibition in normal rats. Our results suggest that RTT-related E/I imbalance in the motor cortex gives rise to excessive synchrony and, consequently, a stereotyped motor code, which may underlie abnormal motor function in RTT.

## Introduction

Orchestration of complex body movements requires some, but not too much, coordination among neurons in motor cortex; neither total synchrony nor total asynchrony will do. Indeed, it is well understood theoretically that excessive correlations can limit the information capacity of any neural code^1–3^ - if all neurons are perfectly synchronized, then different neurons cannot encode different functions. Precisely how much and what kind of synchrony is best for motor coding and how motor function depends on changes in synchrony is poorly understood. Previous studies have shown that significant synchrony and correlations exist among neurons in motor cortex^4–7^. Is this coordinated activity beneficial or is it actually limiting the motor code? Recent experiments have shown that correlations can limit motor coding^8^ and that the neurons with strongest correlations in M1 are weakly related to body movements^7^. Nonetheless, most studies of synchrony or correlations among neurons in motor cortex have focused on how such coordinated activity may play a beneficial role in motor coding^9–12^. For instance, particular groups of synchronized neurons seem to send control signals to particular muscle groups^9,10^ and propagation of correlated firing contributes to motor planning^12^. These findings suggest that correlated activity among specific subsets of neurons encode specific motor functions. Thus, it stands to reason that if this synchrony became less selective and more widespread, then the motor code would lose specificity and motor function could be compromised.

Here we explored this possibility in primary motor cortex of rats under two conditions. First, we studied a transgenic rat model of Rett syndrome (RTT), which has disrupted expression of the MeCP2 gene. Second, we studied normal rats with acutely altered inhibitory neural interactions. Both these cases are associated with abnormal motor behavior and, possibly, abnormal synchrony. Abnormal synchrony is a possibility, because both these cases are linked to an imbalance between excitatory (E) and inhibitory (I) neural interactions, which in turn is likely to result in abnormal synchrony. For instance, many computational models suggest that synchrony is strongly dependent on E/I interactions^13–16^. Likewise, in experiments, pharmacological manipulation of E/I causes changes in synchrony^7,14,17^ and the excessive synchrony that occurs during epileptic seizures is often attributed to an E/I imbalance^18,19^. Similarly, the majority of people with RTT suffer from seizures and many previous studies establish E/I imbalance as a common problem in RTT^20^. MeCP2 dysfunction, which is known to cause RTT, seems to be particularly important in inhibitory neurons^21^. For instance, two studies have shown that disrupting MeCP2 only in specific inhibitory neuron types can recapitulate the effects of brain-wide disruption of MeCP2^22,23^. However, whether the E/I imbalance favors E or I at the population level seems to vary across different brain regions in RTT. Studies of visual cortex^22^ and hippocampus^24^ suggest that the balance tips towards too much excitation, while studies of somatosensory cortex^25,26^ and a brain-wide study of Fos expression^27^ suggest that forebrain areas, including motor cortex, are tipped towards excessive inhibition. These facts motivated our choice to study pharmacological disruption of inhibition here. While it is clear that E/I imbalance is important in RTT, it is much less clear how it manifests at the level of dynamics and correlations in the neural activity that is responsible for coordinating body movements. Thus, in addition to pursuing the general questions relating synchrony and motor function discussed above, a second goal of our work was to improve understanding of motor dysfunction due to MeCP2 disruption.

Taken together, these facts led us to the following experimental questions. How are abnormal body movements and motor coding related to abnormal synchrony in MeCP2-disrupted motor cortex? Are abnormalities in the MeCP2-disrupted motor system consistent with excessive inhibition in motor cortex? Our work here shows that both MeCP2 disruption and excessive inhibition lead to excessive synchrony in motor cortex. The degree of synchrony is correlated with the degree of abnormality of behavior and motor coding. Our findings suggest that RTT-related motor dysfunction may be due, in part, to excessive synchrony and inhibition in motor cortex.

## Results

We performed simultaneous measurements of body movements and spiking activity of many single neurons in motor cortex of rats (Fig. 1a,b). We compared four different experimental groups: normal rats (rattus norvegicus, *n* = 3, weights, Sprague Dawley, Harlan Labs, TX, USA), normal rats with pharmacologically altered inhibition, transgenic RTT rats (*n* = 4, weights, HET KO, SD-Mecp2tm1sage, Horizon Labs, MO, USA), and RTT rats with pharmacologically altered inhibition. The rat model of RTT we study here has been shown to recapitulate important dysfunctions and behaviors found in RTT humans including impaired motor functions^28–30^. Our goal was to quantitatively assess differences in three properties of these four experimental groups: body movements, neural activity, and the neuron-to-body relationship.

**Figure 1.**
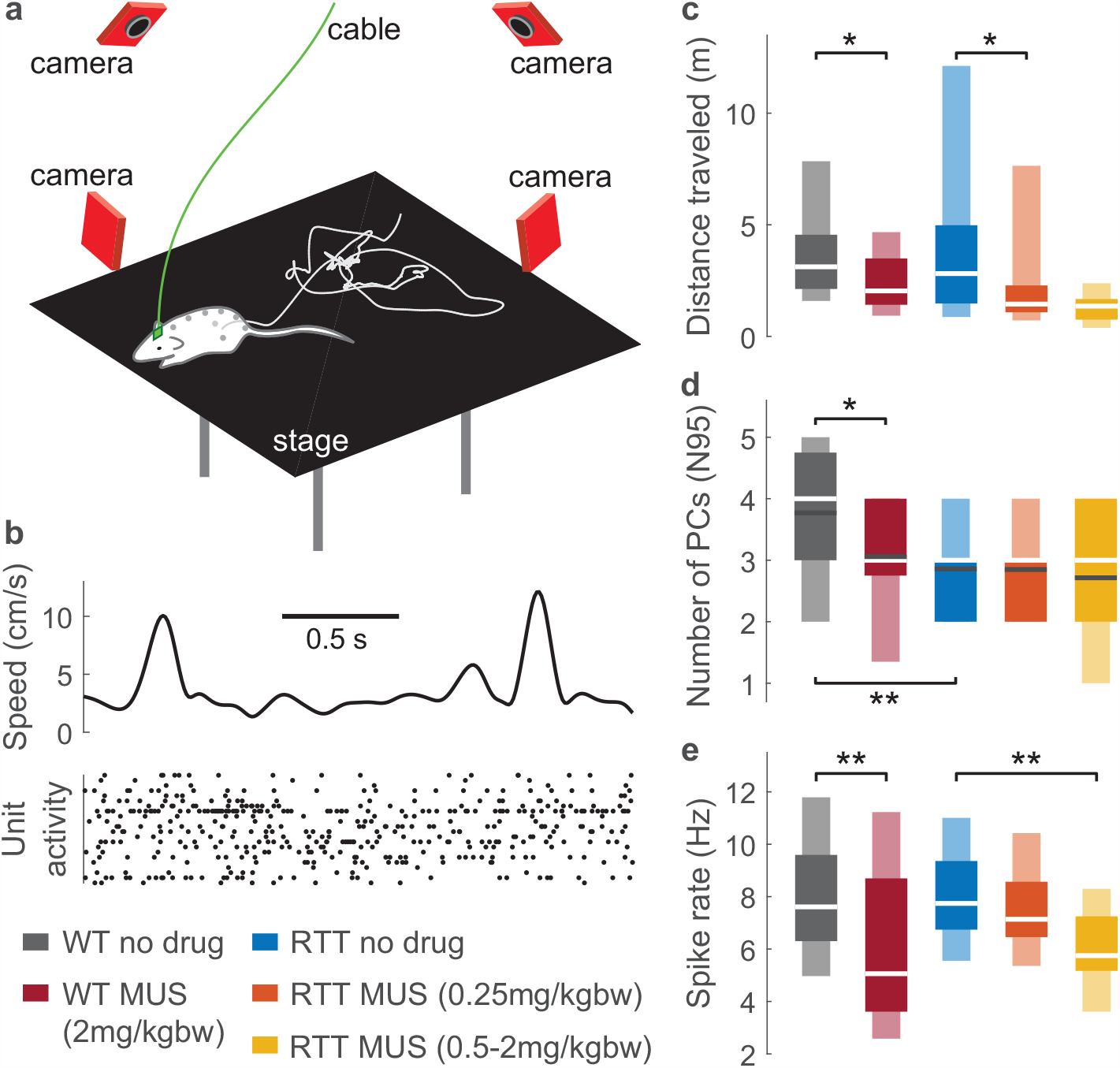
Reduced behavioral complexity due to MeCP2 disruption and excessive inhibition in motor cortex. **a**, An example movement path (white line) of a rat, obtained by tracking the positions of eight beads attached to the rat. **b**, Body speed and single unit neural activity were recorded simultaneously. **c**, Both wild-type (WT) rats and RTT rats moved less due to enhanced inhibition (WT no drug vs. MUS: *p* = 0.022; RTT no drug vs. MUS low dose: *p* = 0.017). **d**, Complexity of movements is reduced for RTT rats compared to WT rats (*p <* 0.001). The reduced complexity is also found in WT rats with enhanced inhibition (*p* = 0.018). **e**, Spike rate is reduced for enhanced inhibition (WT no drug vs. MUS: *p* = 0.010; RTT no drug vs. MUS high dose: *p <* 0.001), but is not different between WT and RTT rats (*p* = 0.978). In **c**-**e**, asterisks indicate t-test significance: \**p <* 0.05, \*\**p <* 0.01. Dark and light boxes delineate 0.25-0.75 and 0.05-0.95 quantiles, respectively. White lines mark median. Black lines mark mean in **d**.

During each 30 minute recording session (*n* = 193 sessions in total), the rats behaved freely - e.g. walking, grooming, and changing posture - on a 30 cm *×* 30 cm platform inside a dark enclosure. To capture body movement, we recorded the three-dimensional positions of eight reflective beads positioned along the neck, back, rear hips, and the base of the tail of each rat, using a infrared multi-camera motion tracking system (Optitrack Flex: V100R2) with millimeter spatial resolution and 10 ms temporal resolution. We analyzed the speeds of the tracking bead and found that administering GABA agonist (muscimol, see methods) resulted in reduced animal motility (Fig. 1c). Although RTT rats were more variable, they did not exhibit significantly different amount of movement on average compared with normal rats (Fig. 1c). However, a more subtle analysis revealed that RTT rats exhibited less complex (i.e. lower dimensional) body movement compared to normal rats (Fig. 1d). We assessed complexity of movements based on principle component analysis of the 8-dimensional bead speed data, considering only times when the animal was active to avoid confounding movement complexity with general motivation to move (see methods). We counted how many principal components were needed to account for 95 percent of the variance of bead speed fluctuations; we called this count N95. We found that normal rats moved with more complex, higher dimensional body motion, requiring significantly more principal components (greater N95) than RTT rats. In addition, we found that, when muscimol was applied, the average N95 of normal rats dropped to a similar level as RTT rats.

Next we analyzed our neural recordings seeking correlates of the reduced complexity in behavior induced by MeCP2 disruption and excessive inhibition. Neural activity was recorded (Cerebus, Blackrock Microsystems) with a 32 channel microelectrode array chronically implanted in deep layers of primary motor cortex (1300*µ*m from the pia). The electrode array sampled from locations associated with diverse muscle groups including hips, neck, whiskers, and more^31^. After spike sorting (Kilosort^32^), we obtained 5197 single units in total (*n* = 1354 from normal rats, *n* = 3843 from RTT rats, average *n* = 27 units per recording). We first compared spike rates across the different experimental groups. As expected, muscimol resulted in decreased spike rates for both normal and RTT rats (Fig. 1e). But, the difference in spike rates between normal and RTT rats was not significant (Fig. 1e).

Next, we examined correlations in the spiking activity. For each recording, we computed the correlation of every pair of simultaneously recorded neurons (Fig. 2a) and then averaged these pairwise correlations across all pairs (Fig. 2b) to obtain a single number, called ‘synchrony’ in Fig. 2c (more detail in Methods). Although synchrony varied greatly across recordings, we found a significant increase in synchrony for the RTT rats compared to the normal rats. Similarly, application of muscimol resulted in a large increase in synchrony for both normal and RTT rats. This result is somewhat surprising considering that stronger inhibition is often associated with reduced synchrony in theory^13,33^ and GABA agonists can result in reduced synchrony^14^, but our finding is consistent with a recent study that applied muscimol in motor cortex of awake rats^7^.

**Figure 2.**
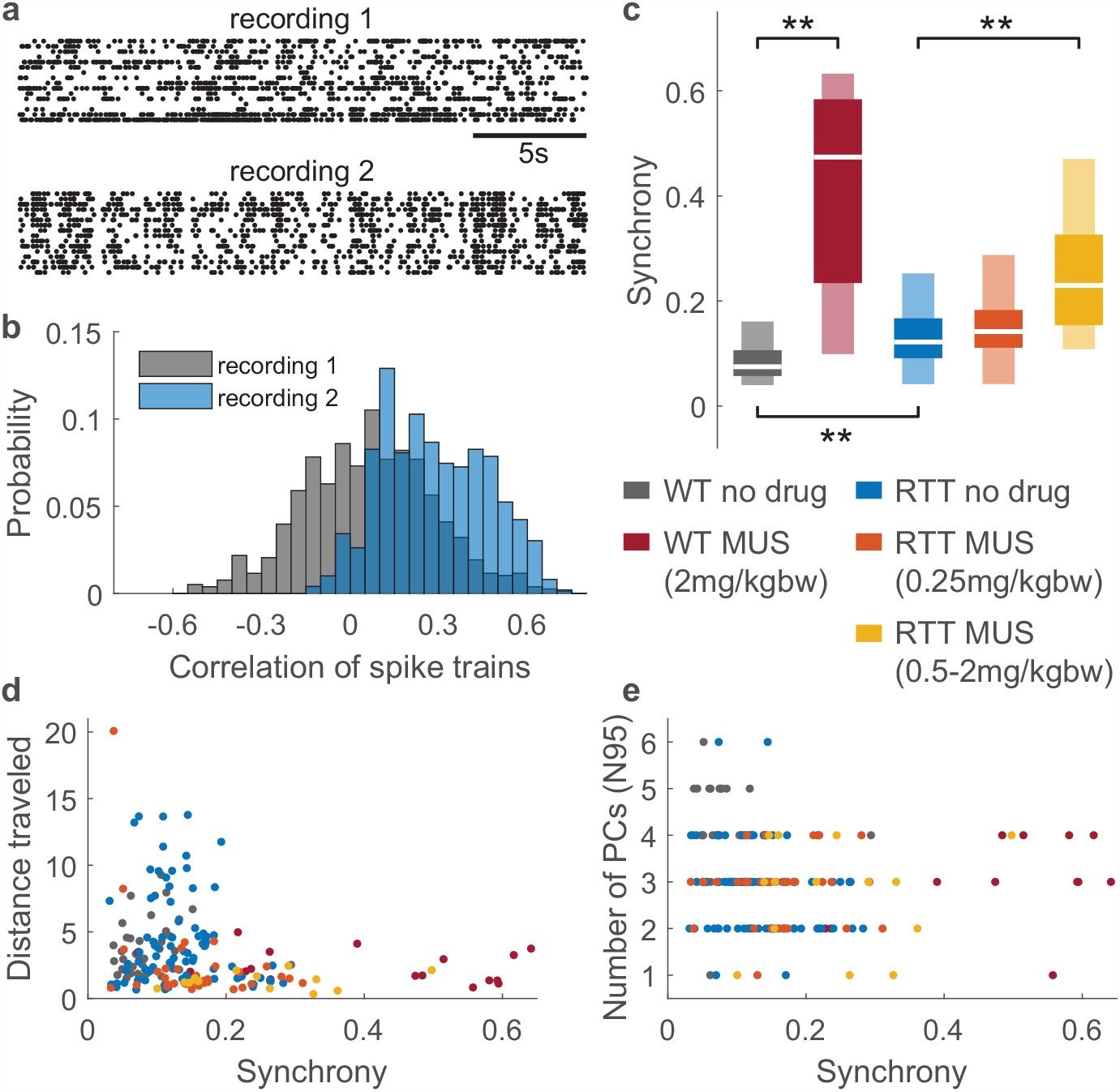
Increased synchrony due to MeCP2 and excessive inhibition correlates to reduced behavioral complexity. **a**, Example spike rasters from a WT no drug recording (recording 1) and a RTT no drug recording (recording 2) show different level of correlations. **b**, Distributions of pairwise correlations of spike trains reveal larger correlations on average in recording 2 compared to recording 1. We defined the average across all pairs as ‘synchrony’. **c**, Summary of all recordings shows increased synchrony in RTT rats compared to WT rats (*p <* 0.001). Similarly, increasing inhibition results in increased synchrony (WT no drug vs. MUS: *p <* 0.001; RTT no drug vs. MUS high dose: *p <* 0.001). Asterisks indicate t-test significance: \*\**p <* 0.01. Dark and light boxes delineate 0.25-0.75 and 0.05-0.95 quantiles, respectively. White lines mark median. **d**, Mobility is negatively correlated with synchrony (Pearson *ρ* = *−* 0.21, *p* = 0.003). **e**, Complexity of movements is negatively correlated with synchrony (Spearman *ρ* = *−* 0.22, *p* = 0.002). Color represents different groups in **d** and **e**.

These results suggest that the increased synchrony we observed in RTT rats and for enhanced inhibition could be responsible for reduced behavioral complexity. Consistent with this possibility, we found that synchrony was negatively correlated with distance traveled (Fig. 2d) and behavioral dimensionality N95 (Fig. 2e).

The synchrony results presented in Fig. 2 provide a limited view of the relationships among neurons. In particular, such average pairwise correlations assess instantaneous relationships and miss any temporally structured interactions among neurons, which may be important if spike timing matters for motor coding. To better account for such temporal relationships, we next examined spike-triggered average histogram waveforms of population activity for each neuron (Fig. 3a); hereafter we refer to these as intracortical interaction (ICI) waveforms. Similar to previous studies of population coupling^7,34,35,^ we define the population activity as the spike count time series of all recorded neurons, excluding the trigger neuron (the one whose spike times are used to do the spike-triggered average). The shape of an ICI waveform reveals whether and how the trigger neuron leads or lags the activity of the network in which it is embedded. A flat line in the ICI waveform would indicate a neuron that fires independently of the population. After obtaining an ICI waveform for each single neuron, we then compared ICI waveforms across neurons. We found that for normal rats, ICI waveforms were diverse; some neurons lead, others lag the population; some neurons had sharply peaked ICI waveforms, others had broader peaks (Fig. 3a, gray). In contrast, in the RTT rats, neurons tended to have stereotyped ICI waveforms (Fig. 3a). This means that each neuron tends to participate with the population in the same way in RTT M1. We quantified this cortical stereotypy by calculating correlations between all pairs of ICI waveforms (Fig. 3b) for each recording and then averaging across all pairs, to obtain a single ICI stereotypy number for each recording. Comparing across our experimental groups, we found that ICI stereotypy was much greater in RTT rats than in normal rats (Fig. 3c). Similarly, we found that application of muscimol resulted in greatly increased ICI stereotypy for both normal and RTT rats (Fig. 3c). We conclude that diversity of temporal relationships among neurons in M1 is compromised in MeCP2 deficient rats in a way that is consistent with an increase in inhibition in M1.

**Figure 3.**
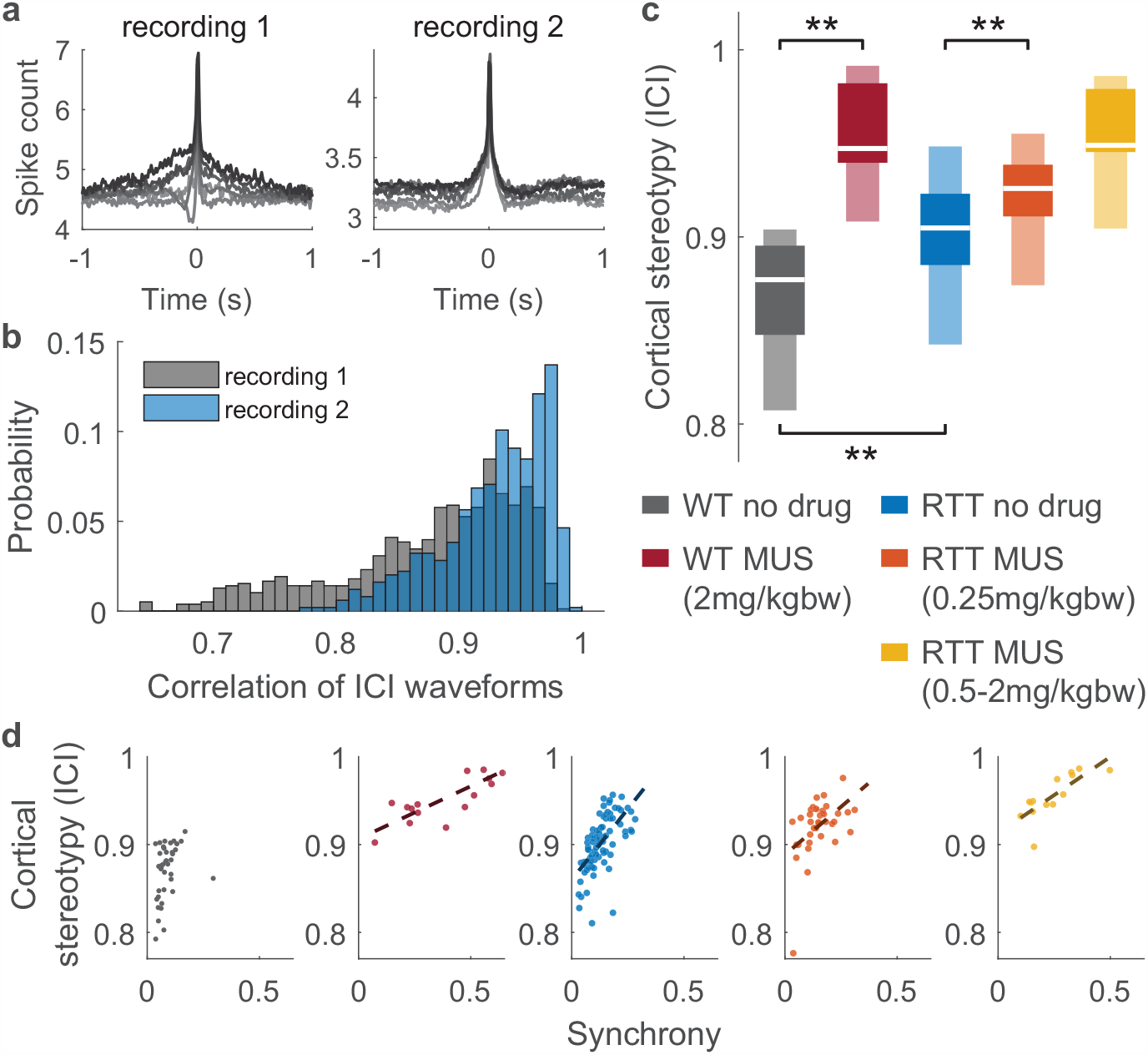
Stereotyped interactions among M1 neurons due to MeCP2 disruption and excessive inhibition. **a**, Spike-triggered average population acitivty (i.e. ICI waveform) for six example neurons from a WT rat (recording 1) and from a RTT rat (recording 2). Stereotyped ICI waveforms in recording 2 indicate that intracortical interactions are less diverse than in recording 1. **b**, ICI waveforms were compared for each pair of neurons. Distributions of pairwise correlations of ICI waveforms reveal greater diversity of waveforms in recording 1 compared to recording 2. We defined the average across all pairs as ‘cortical stereotypy’. **c** Summary of all recordings shows that RTT rats have greater cortical stereotypy than WT rats (*p <* 0.001). Increased cortical stereotypy is also caused by enhanced inhibition (WT no drug vs. MUS: *p <* 0.001; RTT no drug vs. MUS low dose: *p* = 0.002). Asterisks indicate t-test significance: \*\**p <* 0.01. Dark and light boxes delineate 0.25-0.75 and 0.05-0.95 quantiles, respectively. White lines mark median. **d**, Excessive synchrony links to increased cortical stereotypy for RTT rats and enhanced inhibition. Cortical stereotypy and synchrony are nearly uncorrelated for WT rats (*ρ* = 0.32, *p* = 0.046), but strongly correlated in RTT rats (*ρ* = 0.61, *p <* 0.001) and enhanced inhibition (WT MUS: *ρ* = 0.83, *p <* 0.001; RTT MUS low dose: *ρ* = 0.47, *p* = 0.003; RTT MUS high dose: *ρ* = 0.78, *p* = 0.001). Color represents different groups. Dashed lines show the best linear fit when *p <* 0.01.

Given the conceptual similarity between synchrony and ICI stereotypy, one might expect that these two quantities are directly linked. However, we found that synchrony and ICI stereotypy were nearly uncorrelated for normal rats (Fig. 3d). In contrast, for RTT rats and for normal rats with increased inhibition, synchrony is strongly correlated with ICI stereotypy (Fig. 3d). We conclude that, even though synchrony can be high in normal rats, this ‘normal synchrony’ does not come with stereotyped intracortical interactions.

So far, we have shown reduced complexity of neural activity (i.e. higher synchrony and greater ICI stereotypy) and reduced complexity of behavior (lower N95) in RTT rats compared to normal rats. Since neurons in M1 influence the body movements of these animals, it stands to reason that the relationships between neurons and body movements, i.e. the motor code, might also be compromised and less complex in RTT rats. Alternatively, it could be that higher-level decision making or movement planning circuits are altered in RTT rats, and that motor coding remains intact. To begin to distinguish these possibilities, we next measured stereotypy of body-cortex interactions (BCI). Similar to our approach with ICI stereotypy, we calculated spike-triggered average waveforms of body speed -termed BCI waveforms (Fig. 4a, more details in methods). We obtained one BCI waveform for each neuron. A complex motor code would manifest with different neurons firing in relation to different aspects of body movement, i.e. diverse BCI waveforms across neurons. For each recording session, we quantified stereotypy of BCI waveforms by calculating the average pairwise correlation of all BCI waveforms (Fig. 4b). Similar to our ICI stereotypy results, we found that BCI waveforms were very diverse in normal rats - BCI stereotypy was low (Fig. 4c). In RTT rats, BCI stereotypy was highly variable and slightly increased on average compared to normal rats, but not significantly. However, the extreme values of BCI stereotypy (say, above 0.7) that we saw in some RTT rats never occurred for normal rats. Increased inhibition resulted in dramatic increases in BCI stereotypy for both normal and RTT rats. How does BCI stereotypy relate to synchrony? Similar to ICI stereotypy, we found that BCI stereotypy was strongly correlated with synchrony for RTT rats and for conditions with excessive inhibition (Fig. 4d). Moreover, for normal rats, BCI stereotypy was not significantly correlated with synchrony. We conclude that the synchrony that occurs naturally among healthy M1 neurons does not entail a low-dimensional motor code like it does in RTT rats and for excess inhibition.

**Figure 4.**
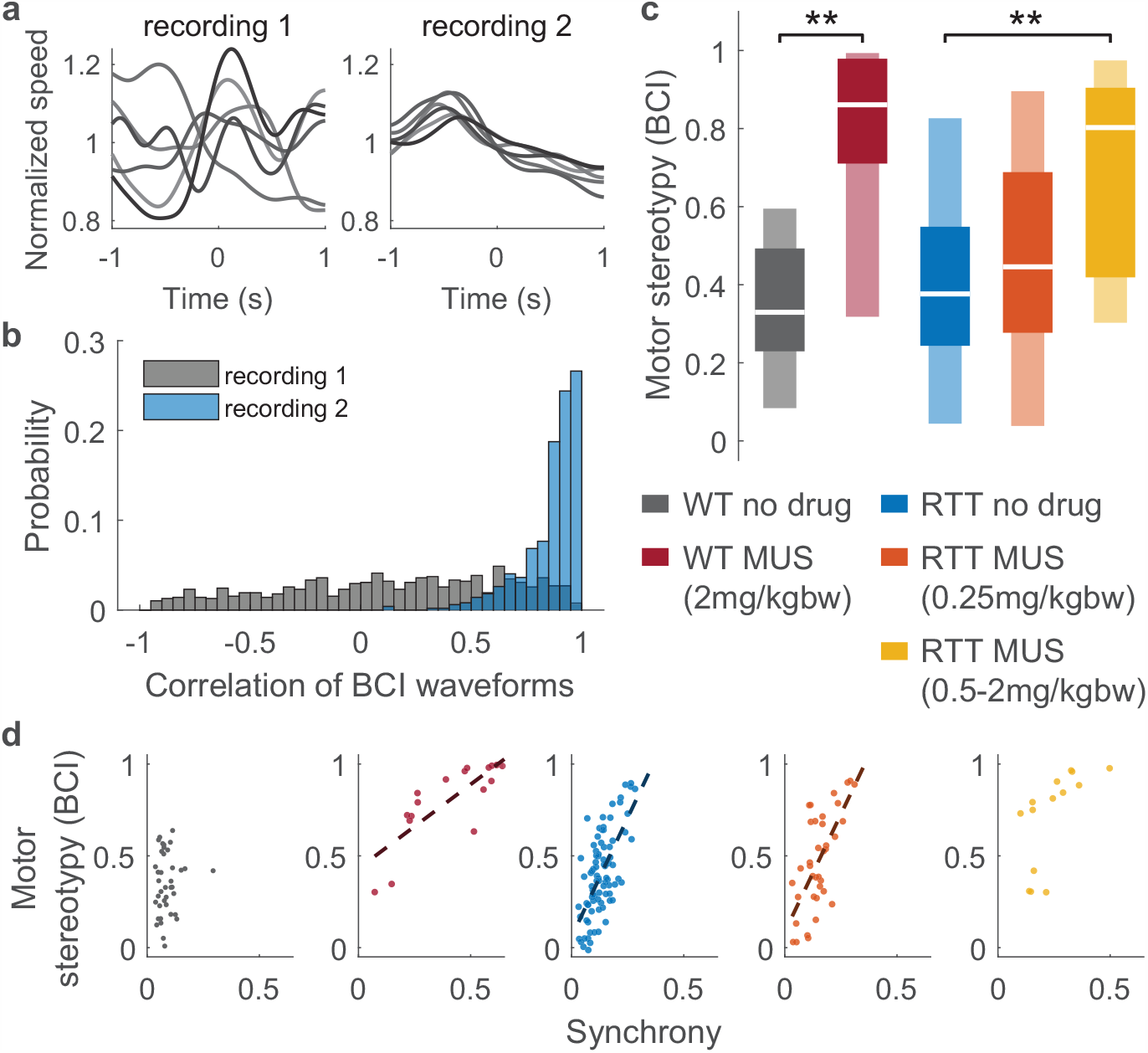
Stereotyped body-cortex interaction due to MeCP2 disruption and excessive inhibition. **a**, Each line (BCI waveform) is a spike-triggered average of body speed from one neuron. These example BCI waveforms are more diverse for the WT rat (recording 1) than for the RTT rat (recording 2). **b**, Summary of BCI waveform similarity (correlation) for all pairs of neuron in recording 1 compared to recording 2. We defined the average across all pairs as motor stereotypy. **c**, Enhanced inhibition caused increased motor stereotypy (WT no drug vs. MUS: *p <* 0.001; RTT no drug vs. MUS high dose *p <* 0.001). Asterisks indicate t-test significance: \*\**p <* 0.01. Dark and light boxes delineate 0.25-0.75 and 0.05-0.95 quantiles, respectively. White lines mark median. **d**, Synchrony correlates with motor stereotypy for RTT rats (*ρ* = 0.66, *p <* 0.001) and enhanced inhibition (WT MUS: *ρ* = 0.80, *p <* 0.001; RTT MUS low dose: *ρ* = 0.72, *p <* 0.001; RTT MUS high dose: *ρ* = 0.65, *p* = 0.012), but not for WT rats (*p* = 0.558). Note that extreme values of motor stereotypy (>0.7) were not observed in normal rats. Color represents different groups. Dash lines show the best linear fit when *p <* 0.01.

## Discussion

Here we have shown that MeCP2 disruption and increased inhibition cause a similar reduction in complexity of the motor system. Compared to normal rats with intact inhibition, this reduced complexity manifested in four ways: body movements were simpler (lower dimensional), neural activity became less diverse (i.e. more synchronous) across the neural population in M1, interactions among neurons in M1 became more stereotyped, and relationships between M1 neurons and body movements became more stereotyped. Returning to the questions we posed at the start, one interpretation of our observations is that MeCP2 disruptions cause an imbalance favoring inhibition in M1, which results in excessive neural synchrony, thereby limiting the information capacity of the motor code; the commands sent to the spinal cord from M1 are less diverse. In this view, the reductions in complexity of behavior and neuron-to-body relationships are a natural consequence of the less complex commands issued from neurons in M1.

However, our more careful look at our results adds nuance to this interpretation. Elevated synchrony in M1 does not always coincide with a general reduction in complexity of the motor system. Indeed, our measurements demonstrate that normal healthy rats exhibit a wide range of synchrony, varying greatly from one recording to another. As shown in Fig. 2c, the 5%-95% range of measured synchrony values in normal rats was 0.04-0.16, which overlaps substantially with the 5%-95% range measured in RTT rats - 0.04-0.25. But, even the highest levels of synchrony we observed in normal healthy rats did not come with dramatic reductions in motor complexity. Considering this fact, a more accurate conclusion from our results would be that enhanced inhibition and MeCP2 disruption result in a dysfunctional type of synchrony, that reduces motor complexity and is different from the synchrony found in healthy M1 circuits. In agreement with this idea, previous work has highlighted the beneficial role of synchrony in M1, “binding” functional groups of neurons which are associated with distinct muscle groups^9,10^ or motor planning^12^. Our measurements of ICI stereotypy suggest that one way in which this unhealthy synchrony differs from normal synchrony lies in the temporal aspects of relationships among neurons. Very high ICI stereotypy (above 0.9), which was not observed in our recordings of normal rats, implies that all neurons have very similar temporal relationships - firing with the same lag or lead relative to the population. In the normal rats, high synchrony could coexist with diverse temporal relationships among neurons.

Application of muscimol in normal rats recapitulated many of our observed differences between RTT rats and normal rats. This observation is consistent with previous studies that point to an imbalance favoring too much inhibition in forebrain areas as a circuit-level problem associated with RTT^25–27^. However, this observation also deserves more careful attention. Not all aspects of our measurements of RTT rats paralleled the effects of muscimol application in normal rats. For example, spike rates were not significantly different between normal and RTT rats. This suggests that if excessive inhibition is indeed an important aspect of RTT, then there are also compensatory mechanisms to keep spike rates in a normal range. Such compensatory mechanisms are a well known challenge of studying long term E/I imbalance in neural disorders^20^. To further explore the role of inhibition in our RTT rat model, we performed additional experiments in which we reduced inhibition by applying GABA antagonists. Our original motivation for this was the possibility of rescuing normal motor function in RTT rats. However, we found that partially blocking inhibition did not recover normal motor function. Perhaps consistent with compensatory mechanisms, we found that RTT rats were more sensitive to reduced inhibition than normal rats, but we did not find a return to normal motor function (Supp Figs. S1 and S2).

Our work highlights the complex role of synchrony in motor system function and dysfunction. We show that MeCP2 disruption can lead to excessive synchrony and a collapse of complexity in the relationships among M1 neurons and the relationships between M1 neurons and the body. Our findings suggest that stereotypy at the level of motor coding may play an important role in the stereotyped body movements of Rett syndrome.

## Methods

### Animals

All procedures followed the Guide for the Care and Use of Laboratory Animals of the National Institutes of Health and were approved by University of Arkansas Institutional Animal Care and Use Committee (protocol #14048). We studied normal Sprague–Dawley male rats (*n* = 3, Harlan Labs, TX, USA) and transgenic MeCP2 knockout female rats (*n* = 4, HET KO, SD-Mecp2^*tm*1*sage*^, Horizon Lab, MO, USA). The raw data from the normal rats was collected and first reported in our previous study^7^, but reanalyzed here. The RTT rats have a 71 base pair deletion in Exon 4, and are maintained by breeding heterozygous females with wild type males, both with Sprague Dawley backgrounds. This animal model has been shown in other studies to recapitulate important dysfunctions and behaviors found in RTT humans including breathing abnormalities, unusual social interactions, exaggerated response to auditory stimuli, reduced gross locomotion, weak grip, and shortened lifespan^28,29^. In addition, these rats have also been shown to manifest many of the same behavioral abnormalities found in some RTT mouse models including stunted body growth, maloccluded teeth, and reduced interest in social novelty^28^.

### Pharmacology

The rats were given either 2 ml sterile saline or muscimol diluted in saline solution through intraperitoneal (IP) injection 50 minutes before every recording. Muscimol is a GABA_A_ agonist that increases the strength of inhibitory signaling^36^. For normal rats, the dose for muscimol is 2 mg/kg body weight; for RTT rats, we applied lower dose varying from 0.25 to 2 mg/kg body weight because these animals seemed to be more sensitive to altered inhibition than normal rats. On each recording day, we performed one no-drug recording first and one muscimol recording after at least 1 hour for each rat.

### Electrophysiology

Microelectrode arrays were chronically implanted 1300 um deep in a 2 mm*×* 2 mm craniotomy with the center located 0.5 mm posterior to bregma and 2 mm lateral from midline. Thus, the recorded neurons were located in deep layers of the primary motor cortex and at positions associated with a wide range of body movements^31^. For the normal rat recordings, we used microelectrode arrays (A8×4–2mm-200–200–413-CM32, Neuronexus); for RTT rats, we used microelectrode array (Buzsaki32-CM32, Neuronexus) for improved spike sorting^37^. For both groups, the plane of microelectrode arrays was oriented perpendicular to the dorsal surface and parallel to the midline. After implantation surgery, the rats recovered for at least 2 weeks before recordings began. During each 30-minute recording, extracellular voltage fluctuations were recorded with 30 kHz sample rate (Cerebus, Blackrock Microsystems). Signals were digitized by a headstage connected to the electrode, and transmitted by a commutator connected to the recording system. Spike sorting was done with the Kilosort (https://github.com/cortex-lab/KiloSort), a fast and accurate spike sorting algorithm for high-channel count probes^32^. Then, we manually curated the spike sorting results with the graphical user interface Phy (https://github.com/cortex-lab/phy). Criteria for a good unit included clear and distinct waveform shapes, refractory periods in auto-correlograms, stability in amplitudes, and distinct principal components in feature space.

### Motion tracking

As in our previous work^7^, body movement was recorded with a infra-red nine-camera motion tracking system (OptiTrack Flex:V100R2), where the three-dimensional coordinate of eight reflective beads (MCP1125, Naturalpoint, 3 mm diameter) temporarily adhered along the spine from neck to tail and on each lateral side of rear hips. The tracking system has 10 ms time resolution and submillimeter spatial resolution. The recordings took place in a dark enclosed space. During a 30-minute recording, the rats were allowed to freely move on a 30 cm *×* 30 cm platform placed at the center of the recording space. The lightweight cable is attached to the ceiling and the length is carefully measured so that it does not impede the free movement of the rats. Each rat went through three acclimatization sessions before recording with the same setup to avoid stress and anxiety. After recordings were completed, the tracking trajectories were manually corrected with the software, Motive (https://optitrack.com/software/motive) and smoothed by a 5 Hz low-pass filter. The speed of center of mass and beads were then obtained by calculating differentiated positions.

### Body tracking data analysis

We used the distance traveled by the rat during the recording to represent the general motility (Figs. 1c and 2d). The distance traveled was calculated for each recording as the cumulative distance traveled by the center of mass of the tracking beads. Complexity of movements (Figs. 1d and 2e) was assessed by principle component analysis on the speed of 8 beads, excluding time periods when the rats were at rest for more than 1.5 s. We defined the animal to be ‘at rest’ if it met two conditions: 1) speed of center of mass less than 0.8 cm/s; 2) speed of each bead less than 1 cm/s. Brief periods of motion, shorter than 0.5 s, preceded and followed by rest were considered rest. After excluding rest periods and applying principle component analysis, we counted the number of principal components that explains 95 percent of variance, defined as N95.

### Spike data analysis

Spike rate of a recording was obtained by the average spike rate across all units during the recording (Fig. 1e). ‘Synchrony’ (Figs. 2c-e, 3d, and 4d) was defined as the average of pairwise correlations (Fig. 2b) of spike count time series across all pairs of neurons. The spike count time series were calculated for each neuron using 1 second time bin.

Cortical stereotypy (Fig. 3c-d) was defined to capture stereotypy in intracortical interactions (ICI). For each neuron (trigger neuron), we counted the number of spikes from the population (with 10 ms time bin) in a ± 1 second time window centered on the spike times of the trigger neuron, excluding the spikes from itself. We then averaged these spike-triggered spike count waveforms across all spikes from the trigger neuron to obtain the ICI waveforms (Fig. 3a). Cortical stereotypy was calculated by averaging pairwise correlations of ICI waveforms across all pairs of units as a single number for each recording (Fig. 3b).

Motor stereotypy (Fig. 4c-d) was defined to capture stereotypy in body-cortex interactions (BCI). Similar to ICI waveform, we obtained a BCI waveform for each neuron (the trigger neuron) by averaging the speed of center of mass in a 1± second time window centered on the spike times of the trigger neuron (Fig. 4a). The waveform was then smoothed by a 1.5 Hz low pass filter and normalized by its mean. Motor stereotypy was defined as the average of pairwise correlations of BCI waveforms across all pairs of units (Fig. 4b) to obtain a single number for each recording.

### Statistics

We examined the statistical significance of the difference between two groups using p-value of t-tests. The p-value represents the probability of accepting the null hypothesis that the means of two groups are not different. Pearson’s correlation coefficient and its corresponding p-value were used to test the correlations between two quantities in Figs 2d, 3d, and 4d. Spearman’s correlation coefficient and its corresponding p-value were used for Fig. 2e. For both types of correlation, the p-value represents the null hypothesis that the two quantities are uncorrelated. We obtained *n* = 193 recordings with at least 8 good units in total (*n* = 39 for wild-type no drug; *n* = 17 for wild-type muscimol; *n* = 86 for RTT no drug; *n* = 37 for RTT muscimol 0.25 mg/kgbw; *n* = 14 for RTT muscimol 0.5-2 mg/kgbw). All these recordings were included in our analysis of spike rate, synchrony, and cortical stereotypy. For analysis of distance traveled, complexity of movements (N95), and motor stereotypy, *n* = 5 recordings were excluded due to absence of motion tracking data (*n* = 1 in RTT no drug and *n* = 4 in RTT muscimol 0.25 mg/kgbw).

## Acknowledgements

J.L., W.S., and S.H.G. were funded by National Institutes of Health (NINDS) grant 1R15NS116742-01. W.S., P.A.K., and S.H.G. were funded by a Foundational Questions Institute grant FQXi-RFP3–1343. W.S., P.A.K. and S.H.G. were funded by Arkansas Biosciences Institute.

## Author contributions statement

W.L.S., J.L., P.A.K., and S.H.G. conceived the experiments; J.L., P.A.K., and S.H.G. conducted the experiment; J.L. and W.L.S. analysed the results and wrote the manuscript. All authors reviewed the manuscript.

## Additional information

The authors declare no competing interests.

---

**Supplementary Figure S1.**
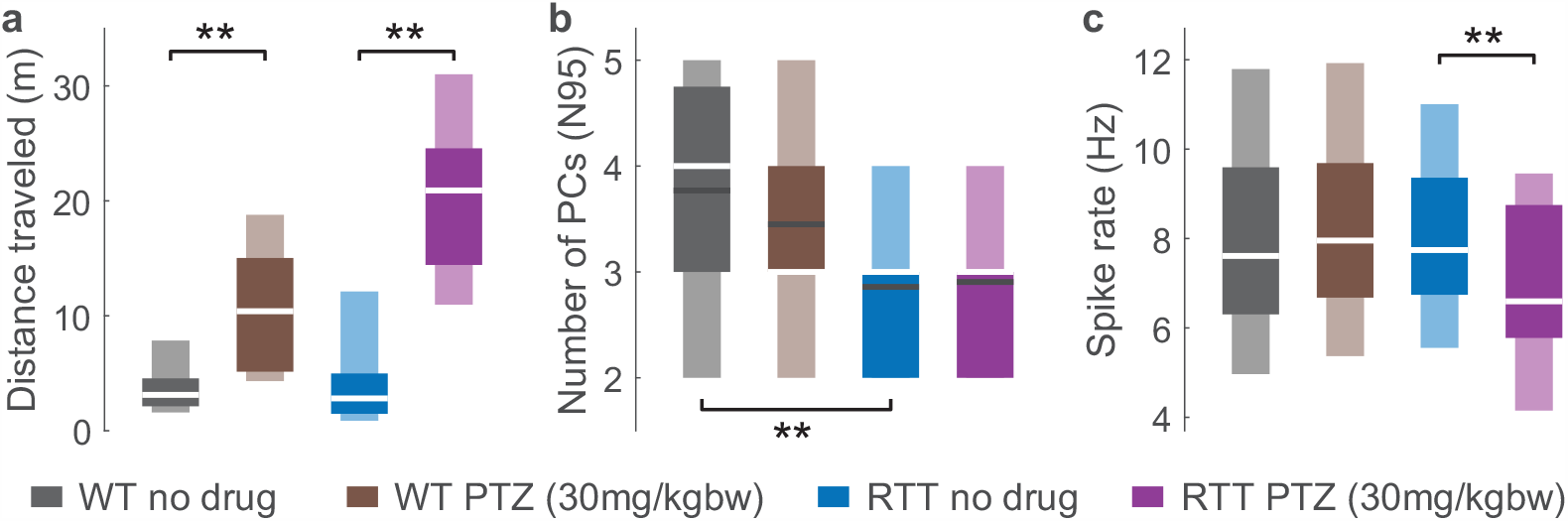
Mobility, complexity of movements, and spike rate for normal rats and RTT rats with reduced inhibition. Pentylenetetrazol (PTZ) was applied as an antagonist of GABA_A_ receptor that decreases the strength of inhibitory signaling. It was diluted in 2 ml saline resolution and induced via IP injection. Animals, electrophysiology, motion tracking, and data analysis remain the same as in method. **a**, Wild-type (WT) rats and RTT rats moved much more due to reduced inhibition (WT no drug vs. PTZ: *p <* 0.001; RTT no drug vs. PTZ: *p <* 0.001). The increase for RTT rats were larger than WT rats. **b**, No significant difference in complexity of movements is found for reduced inhibition (WT no drug vs. PTZ: *p* = 0.246; RTT no drug vs. PTZ: *p* = 0.774). **c**, Spike rate is not different in WT rats with reduced inhibition (*p* = 0.827), but is reduced in RTT rats with reduced inhibition (*p* = 0.001). Asterisks indicate t-test significance: \*\**p <* 0.01. Dark and light boxes delineate 0.25-0.75 and 0.05-0.95 quantiles, respectively. White lines mark median. Black lines mark mean in **b**.

**Supplementary Figure S2.**
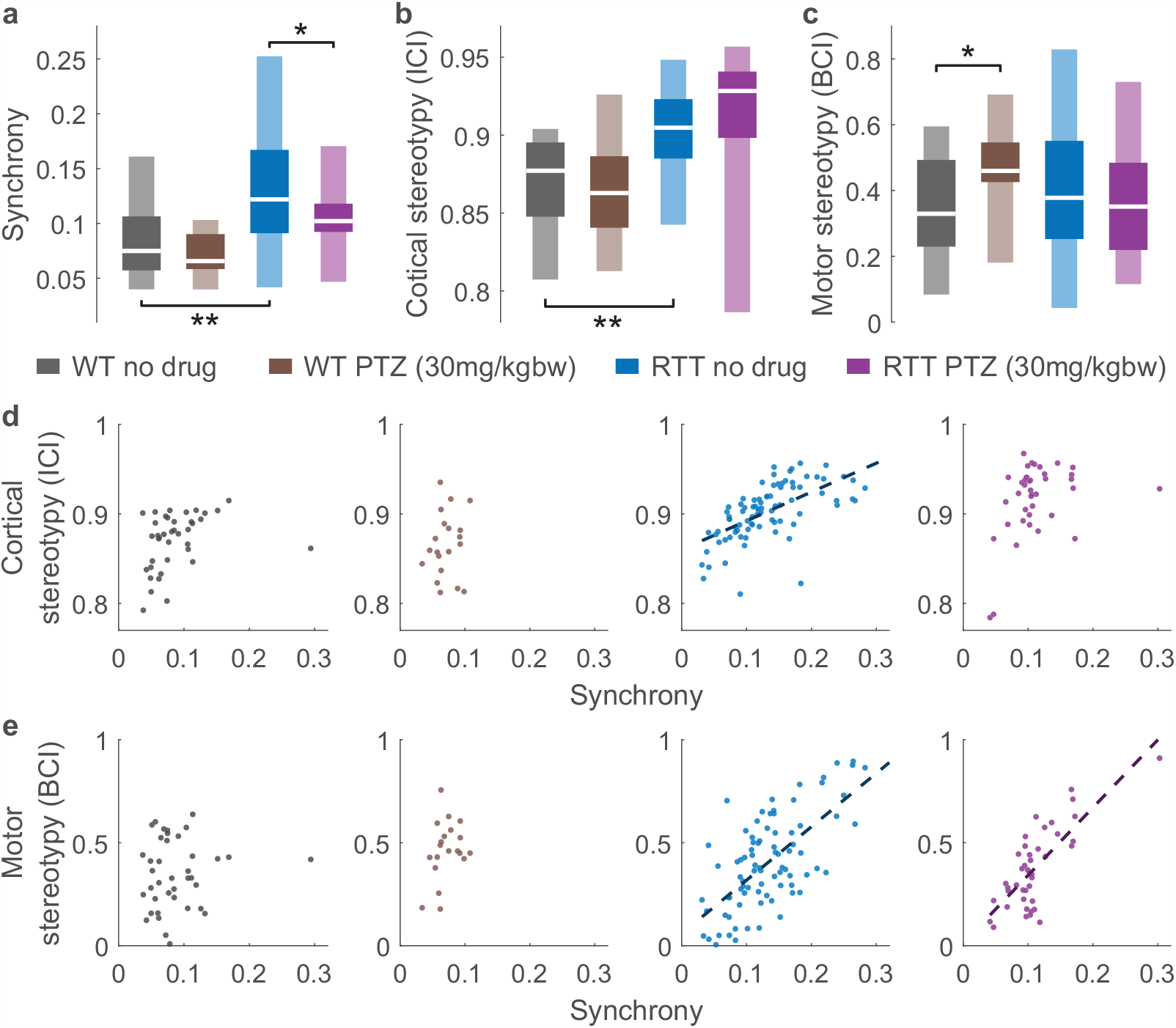
Synchrony, cortical stereotypy, motor stereotypy, and their correlations for normal rats and RTT rats with reduced inhibition. **a**, Synchrony is not significantly different in reduced inhibition for WT rats (*p* = 0.146), but decreases in RTT rats with reduced inhibition (*p* = 0.031). **b**, No significant difference is found in incortical stereotypy for reduced inhibition (WT no drug vs. PTZ: *p* = 0.587; RTT no drug vs. PTZ: *p* = 0.111). **c**, Reduced inhibition caused increased in motor stereotypy for WT rats (*p* = 0.010), but not for RTT rats (*p* = 0.475). In **a**-**c**, asterisks indicate t-test significance: \**p <* 0.05, \*\**p <* 0.01. Dark and light boxes delineate 0.25-0.75 and 0.05-0.95 quantiles, respectively. White lines mark median. **d**, Cortical stereotypy and synchrony stayed uncorrelated for WT rats with reduced inhibition (*p* = 0.552). For RTT rats, the correlation of cortical stereotypy and synchrony was weaken with reduced inhibition (*ρ* = 0.38, *p* = 0.012). **e**, Motor stereotypy and synchrony stayed uncorrelated for WT rats with reduced inhibition (*p* = 0.160), and stayed correlated for RTT rats with reduced inhibition (*ρ* = 0.76, *p <* 0.001). In **d**-**e**, color represents different groups. Dash lines show the best linear fit when *p <* 0.01.

